# Neuroticism as a predictor of frailty in old age: a genetically informative approach

**DOI:** 10.1101/527135

**Authors:** Hilda Bjork Danielsdottir, Juulia Jylhävä, Sara Hägg, Yi Lu, Lucía Colodro-Conde, Nicholas G. Martin, Nancy L. Pedersen, Miriam A. Mosing, Kelli Lehto

## Abstract

**Objective:** Neuroticism is associated with poor health outcomes, but its contribution to the accumulation of health deficits in old age, i.e. frailty, is largely unknown. We aimed to explore associations between neuroticism and frailty cross-sectionally and over up to 29 years, and to investigate the contribution of shared genetic influences.

**Method:** Data were derived from the UK Biobank (UKB, n=502,631), the Australian Over 50’s Study (AO50, n=3,011) and the Swedish Twin Registry (SALT n=23,744, SATSA n=1,637). Associations between neuroticism and the Frailty Index were investigated using regression analysis cross-sectionally in UKB, AO50 and SATSA, and longitudinally in SALT (25-29y follow-up) and SATSA (6 and 23y follow-up). The co-twin control method was applied to explore the contribution of underlying shared familial factors (SALT, SATSA, AO50). Genome-wide polygenic risk scores for neuroticism in all samples were used to further assess whether common genetic variants associated with neuroticism predict frailty.

**Results:** High neuroticism was consistently associated with greater frailty cross-sectionally (adjusted β, 95% confidence intervals in UKB= 0.32, 0.32-0.33; AO50= 0.35, 0.31-0.39; SATSA= 0.33, 0.27-0.39) and longitudinally up to 29 years (SALT= 0.24; 0.22-0.25; SATSA 6y= 0.31, 0.24-0.38; SATSA 23y= 0.16, 0.07-0.25). When controlling for underlying shared genetic and environmental factors the neuroticism-frailty association remained significant, although decreased. Polygenic risk scores for neuroticism significantly predicted frailty in the two larger samples (meta-analyzed total β= 0.06, 0.05-0.06).

**Conclusion:** High neuroticism is associated with the development and course of frailty. Both environmental and genetic influences, including neuroticism-associated genetic variants, contribute to this relationship.

## Introduction

While chronological age is a major determinant of health status, there is substantial diversity in health among older people of the same age (1). One indicator of such variation in health among older individuals is frailty, a clinical condition observed in older people reflecting cumulative decline in various physiological systems (2). Frailty is a strong predictor of mortality (3) and has been linked to numerous other negative health outcomes, such as disability, institutionalization and hospitalization (4).

The causes of frailty are multifactorial, and it is widely accepted that many biological, social and psychological factors are likely involved (5). While most research has focused on biological and physical factors associated with frailty (e.g. body weight) (5), as well as on sociodemographic factors (e.g. older age, female sex, and lower educational level) (6), less is known about how psychological factors could contribute to frailty. Neuroticism, a stable personality trait reflecting a tendency towards emotional instability and negative affect (e.g. depressed mood, worry, fear), has been consistently associated with a wide range of physical and mental health problems such as cardiovascular disease, disrupted immune functioning, asthma, irritable bowel syndrome, atopic eczema, migraine, mood and anxiety disorders, and even increased risk of premature mortality (7-9), potentially also affecting frailty. The association between neuroticism and frailty has been investigated in three previous studies, all suggesting that high neuroticism is associated with higher frailty concurrently and longitudinally, over up to eight years (10-12).

Twin and family studies show moderate heritability of neuroticism, with about 40% of individual differences in the trait attributable to genetic influences (13), potentially contributing to its persistent associations with health problems. Indeed, twin studies also show moderate heritability for many somatic and health-related measures (14), and genetic overlap between neuroticism and some somatic diseases have been detected (15), indicating that neuroticism and health problems could be associated in part because of shared genetic influences.

As a complex phenotype, recent genome-wide association studies (GWAS) have confirmed the complex genetic architecture of neuroticism where many genetic variants with small effects are involved (e.g. 16). By using information from GWAS’s, polygenic risk scores allow testing the contribution of thousands of neuroticism-related common genetic variants in frailty in old age, providing more insight into the potential sources underlying the association.

To date, little is known about the nature of the association between neuroticism and frailty, since study designs used in previous studies do not allow for conclusions about the underlying genetic factors and direction of the relationship. In the present study, we investigated the association between neuroticism and frailty in middle-aged and older adults using four large genetically informative samples. Specifically, we aimed to 1) assess the phenotypic cross-sectional and longitudinal association between neuroticism and frailty, expanding the follow-up time to up to 29 years; 2) assess whether the association between neuroticism and frailty remains after controlling for shared familial influences (shared genes and family environment); and 3) examine whether measured genetic risk for neuroticism contributes to frailty.

## Method

### Data sources / participants

Data were derived from four cohorts of middle aged and older individuals, the UK Biobank (UKB) (17), the Australian Over 50’s study (AO50) (18), and two sub-samples of the Swedish Twin Registry (STR) (19, 20).

The UKB is a large resource of health, lifestyle and genetic data on currently approximately 500,000 individuals aged 39-73 years at recruitment (17). Genotype information was available for 244,070 individuals after exclusions (see details in *Supplementary Material*).

The AO50 is a cross-sectional population-based study of Australian twins older than 50 years of age. The sample consisted of 3,053 individuals aged between 50 and 94 years who answered a mailed-out questionnaire between 1993 and 1995, which included assessments on personality traits, physical and mental health, lifestyle factors, and demographic characteristics (18). Genotype information was available for 1,037 individuals after exclusions.

Screening Across the Lifespan of Twins (SALT) is a cohort study of STR twins born in 1886-1958 (n=44,919) (19). Health and lifestyle data were collected between 1998 and 2002 through a computer-assisted telephone interview. Personality information was available for all SALT participants born 1926-1958 who had completed a mailed-out questionnaire in 1973 (19), resulting in a 25-29-year follow-up between neuroticism and frailty assessments for 24,432 individuals. Genotype information was available for a sub-sample (n=10,712) (20).

The Swedish Adoption/Twin Study of Aging (SATSA) is a longitudinal study of aging spanning over 30 years and includes nine questionnaire-based study waves (21). In 1984 a questionnaire, covering a wide range of health, lifestyle and personality factors, was sent out to all STR twins who were reared apart and a matched sample of twins who were reared together (n=3,838) (21). In the present study we used baseline information from wave 2 (Q2, 1987, n=1,637), and follow-up information from waves 4 (Q4, 1993, n=1,450) and 7 (Q7, 2010, n=568), providing follow-up data over six (wave 4) and 23 years (wave 7). In total, there are 929 individuals with both baseline and six-year follow-up information and 191 individuals with both baseline and 23-year follow-up information. Waves 2, 4 and 7 were selected based on data availability and to maximize sample size in longitudinal analyses. Sample overlap between SATSA and SALT was removed from all SALT analyses. Genotype information was available for 637 individuals.

### Neuroticism assessment

In the UKB and AO50, neuroticism was measured with a 12-item version of the Neuroticism scale from the Eysenck Personality Questionnaire-Revised (EPQ-R) (22). In the STR, neuroticism was measured with a 9-item version from the EPQ (23). Items were scored as “no” [0] or “yes” [1] and then summed with a higher score indicating higher levels of neuroticism (see *Supplementary Figure 1* for the distribution of neuroticism scores in each sample).

### Frailty Index

The Rockwood Frailty Index (FI) was used to assess frailty and created in each sample following the standard protocol (24). The FI defines frailty as a state caused by the accumulation of health deficits expressed as the proportion of present deficits of the total health deficits considered (24). Here, the derived FIs were based on 49 health deficits in UKB, 44 in SALT, 42 in SATSA, and 40 in AO50, depending on the relevant measures available in each sample. An individual’s FI score constitutes of the number of deficits (for that individual) divided by the total number of deficits composing the FI. Detailed descriptions of the creation and validation of the FIs were reported elsewhere, see (25) for UKB, (26) for SATSA, and (manuscript under submission) for SALT. For a detailed description of the FI in AO50 study see Supplementary Materials. Items overlapping between FIs and the respective EPQ scales were excluded in all four samples (three items in UKB, one item in AO50 and two items in SATSA, see *Supplementary Table 2*) and FIs were recalculated.

### Genotyping

For UKB, imputed genetic data released in 2018 were used. Two custom genotyping arrays were used to cover more than 800,000 markers and were further imputed to Haplotype Reference Consortium (HRC) and UK10K + 1000 Genomes Phase 3 reference panels (27). In AO50, individuals were genotyped using Illumina single-nucleotide polymorphism (SNP) platforms: 317, 370, 610, 660, Core-Exome, PsychChip, Omni2.5 and OmniExpress and were imputed to HRC.1.1. In SALT, genotyping was carried out using the Illumina OmniExpress bead chip, further imputed to Hapmap 2 build 36 reference panel. In SATSA, genotyping was carried out using Illumina PsychArray-24 BeadChip, imputed to 1000 Genomes Phase 3 reference panel. For both STR samples, only one twin from each monozygotic (MZ) pair was directly genotyped and genotypes were later imputed to their co-twin.

### Polygenic risk scores (PRS) for neuroticism

PRS for neuroticism (PRS_N_) were created in the four independent target samples using the results (effect sizes and *p*-values for each SNP) from a GWAS on neuroticism (28), by counting the numbers of risk alleles at independent loci, multiplying with the effect size and summing the values across all investigated SNPs (performed in Plink 1.9 and Plink 2.0). The PRS_N_ were created under eight *p*-value thresholds (*p*_T_), from 5×10^−8^ to 1 in UKB, AO50 and SATSA, and from 0.001 to 1 in SALT and each PRS_N_ was standardized using z-scores. The threshold that explains the highest percentage of variance in neuroticism in each sample was used in the main hypothesis testing.

### Covariates

Variables with a conceptual rationale for being associated both with neuroticism trait scores and FI scores were considered as potential confounders. These included age, sex, education, smoking status, physical activity, and body max index (BMI). In the UKB and AO50, all covariates were measured concurrently with neuroticism and frailty. In SATSA and SALT, all covariates were measured at baseline with the exception of education in SALT, which was concurrent. For more detailed description of covariate assessment, see Supplementary Materials. In analyses including PRS_N_, four to 20 Principal Components (PC) (depending on the target sample size) were included as covariates to account for population stratification.

### Statistical analysis

#### Phenotypic analyses

Multivariable linear regression analyses were used to determine whether neuroticism was phenotypically associated with FI scores cross-sectionally in UKB, AO50 and SATSA, and longitudinally (over 25-29 years in SALT and over six and 23 years in SATSA) (aim 1). In the cross-sectional analyses, we adjusted for age, sex, and educational level (Model 1) and then additionally for smoking status, exercise and BMI (Model 2). In the longitudinal analyses, follow-up FI scores were predicted from baseline neuroticism while adjusting for all covariates (Model 1). To reduce the possibility of reverse causation (i.e. higher baseline frailty influencing neuroticism level), baseline FI score/chronic illness (chronic illness in SALT as baseline frailty could not be derived) was additionally included as a covariate (Model 2). Since the cohorts are composed of related individuals (twins in AO50, SALT and SATSA), dependency between observations due to relatedness was controlled for by using cluster-robust standard error estimator (i.e. the sandwich estimator) on family ID.

#### Co-twin control analyses

Co-twin control analysis was used to examine associations between neuroticism and frailty with regard to familial (genetic and environmental) factors shared within the twin (aim 2). Dizygotic (DZ) twins share on average 50% of their segregating genes and monozygotic (MZ) twins share all their genes, while both MZ and DZ twins share their family environment. If neuroticism’s effect on frailty is beyond familial influences (consistent with a causal hypothesis), we would expect that the twin with higher neuroticism would also be more frail, i.e. the within-pair associations (MZ and DZ) between neuroticism and frailty would be similar in strength to the individual-level association in the whole sample (29). If the association is better explained by shared underlying factors (e.g. genetic factors), the strength of the association would be attenuated within DZ and especially MZ twins (see (29) for further details). Within-pair difference scores for neuroticism and FI were calculated. Linear regression analyses were used to test whether within-pair differences in neuroticism predicted within-pair differences in FI scores, both cross-sectionally (in AO50 and SATSA) and longitudinally (25-29 years in SALT and in SATSA only over six years as the number of full pairs was low after 23 years), adjusting for within-pair differences in education, smoking, exercise, and BMI. As only same-sex twins were included, and twins are by default the same age, possible confounding influences of sex and age were intrinsically adjusted for by the co-twin design.

#### PRS analyses

Multivariable linear regression with PRS_N_ as independent variable adjusting for age, sex and PCs, was used. First, to validate PRS_N_ as a predictor for neuroticism and to determine which PRS_N_ *p*_T_ explained the highest proportion of variance in phenotypic neuroticism in each sample to be used for the main analyses, the difference in *R*^2^ between the full (including PRS_N_) and reduced (including only the covariates) models were compared. The selected PRS_N_ were then regressed on the respective frailty score in each sample to examine whether measured genetic risk for neuroticism predicts frailty (aim 3). The resulting coefficients from each cohort were then combined in a meta-analysis to get an estimate of the overall effect taking into account sample size. Dependency between observations was controlled for by using the cluster-robust standard error estimator on family ID.

Standardized regression coefficients were reported for all regression analyses to enable comparison between models. Statistical analyses were carried out using Stata version 15.

#### Sensitivity analyses

Since neuroticism is strongly correlated with mental health (8), we also created additional FI’s in UKB, AO50 and SATSA cohorts, further removing any mental health items in addition to those part of the neuroticism scale, (i.e. four items in UKB, three items in AO50, and two items in SATSA, see *Supplementary Table 3*) and re-ran the cross-sectional analyses, to assess whether the association between neuroticism and frailty held. Furthermore, item level sensitivity analyses between frailty and neuroticism (i.e. neuroticism items predicting FI score and neuroticism sum score predicting frailty items) were conducted. A sensitivity analysis was also carried out to test the association between PRS_N_ created under all eight *p*_T_s and FI scores.

## Results

Descriptive statistics of baseline characteristics and follow-up FI scores of the four samples are presented in Table 1.

**Table 1.** Baseline characteristics of the samples

|  |  |  |  |  | STR |  |  |  |
| --- | --- | --- | --- | --- | --- | --- | --- | --- |
|  | UKB |  | AO50 |  | SALT |  | SATSA |  |
|  | Mean (SD)<br>/Percent | n | Mean (SD)<br>/Percent | n | Mean (SD)<br>/Percent | n | Mean (SD)<br>/Percent | n |
| **Age (years)** | 56.53 (8.10) | 502.631 | 61.35 (8.60) | 3.011 | 29.14 (8.82) | 23.744 | 62.42 (13.72) | 1.637 |
| **Sex** |  | 502.631 |  | 3.011 |  | 23.744 |  | 1.637 |
| Male | 45.59% |  | 29.53% |  | 47.15% |  | 41.16% |  |
| Female | 54.41% |  | 70.47% |  | 52.85% |  | 58.84% |  |
| **Zygosity** |  |  |  | 3.011 |  | 18.444 |  | 1.563 |
| MZ | na |  | 48.92% |  | 39.79% |  | 34.04% |  |
| DZ | na |  | 50.68% |  | 60.21% |  | 65.96% |  |
| **Neuroticism<sup>a</sup>** | 4.12 (3.27) | 401.665 | 4.00 (3.17) | 2.946 | 2.78 (2.33) | 21.175 | 2.43 (2.22) | 1.575 |
| **Exercise** |  | 502.531 |  | 2.847 |  | 21.123 |  | 1.609 |
| No | 14.82% |  | 11.35% |  | 10.84% |  | 12.06% |  |
| Yes | 85.18% |  | 88.65% |  | 89.16% |  | 87.94% |  |

Table 1 (continued).
|  |  |  |  |  |  |  |  |  |
| --- | --- | --- | --- | --- | --- | --- | --- | --- |
| **Educational level** |  | 407.665 |  | 2.901 |  | 22.100 |  | 1.518 |
| Low | 20.46% |  | 25.75% |  | 73.05% |  | 86.63% |  |
| High | 79.54% |  | 74.25% |  | 26.95% |  | 13.37% |  |
| **Smoking status** |  | 499.679 |  | 2.960 |  | 21.081 |  | 1.460 |
| Non-smoker | 54.75% |  | 50.81% |  | 45.26% |  | 70.68% |  |
| Ex-smoker | 34.64% |  | 35.84% |  | 41.74% |  | 5.96% |  |
| Current smoker | 10.60% |  | 13.34% |  | 13.00% |  | 23.36% |  |
| **BMI** | 27.43 (4.80) | 499.511 | 24.86 (4.02) | 2.973 | 21.74 (2.85) | 20.951 | 24.56 (3.50) | 1.444 |
| **Chronic illness** |  |  |  |  |  | 21.011 |  |  |
| No | na |  | na |  | 85.90% |  | na |  |
| Yes | na |  | na |  | 14.10% |  | na |  |
| **FI score baseline** | 0.12 (0.08) | 500.336 | 0.14 (0.09) | 3.011 | na | na | 0.10 (0.09) | 1.479 |
| **FI score < 10 years follow-up** |  |  |  |  | na | na | 0.09 (0.08) | 1.407 |
| **FI score > 20 years follow-up** |  |  |  |  | 0.12 (0.085) | 23.085 | 0.12 (0.10) | 522 |

*Note.* AO50 = The Australian Over 50's study; BMI = body mass index; DZ = Dizygotic; FI = Frailty index; MZ = Monozygotic; SALT = Screening Across the Lifespan of Twins Study; SATSA = The Swedish Adoption/Twin Study of Aging; SD = Standard Deviation; STR = Swedish Twin Registry; UKB = UK Biobank; <sup>a</sup> - Neuroticism was assessed using different scales.

In all cohorts with cross-sectional data, higher neuroticism was associated with higher FI scores (Table 2) with approximately 0.3 standard deviation (SD) increase in FI scores with each SD increase in neuroticism. Similar results, though somewhat attenuated, were observed when analyses were repeated with FI scores without mental health items (see *Supplementary Table 4*). The longitudinal analyses between neuroticism and frailty in SALT and SATSA showed that high baseline neuroticism was associated with higher frailty measured six, 23, and 25-29 years later (Table 3, *Model 1*). Furthermore, the association remained significant when controlling for baseline chronic illness in SALT as well as frailty in SATSA across six but not 23 years (Table 3, *Model 2*).

**Table 2.** The cross-sectional association between neuroticism and FI scores in UKB, AO50 and SATSA cohorts (Beta and 95% CI).

|  | UKB |  | AO50 |  | SATSA |  |
| --- | --- | --- | --- | --- | --- | --- |
|  | Model 1<sup>a</sup> | Model 2<sup>b</sup> | Model 1 | Model 2 | Model 1 | Model 2 |
|  | n = 324,724 | n = 323,136 | n = 2,849 | n = 2,650 | n = 1,365 | n = 1,318 |
| Neuroticism | 0.33(0.33,0.34) | 0.32(0.32,0.33) | 0.35(0.31,0.39) | 0.35(0.31,0.39) | 0.35(0.29,0.40) | 0.33(0.27,0.39) |
| Age | 0.18(0.18,0.19) | 0.17(0.17,0.18) | 0.23(0.19,0.27) | 0.27(0.23,0.31) | 0.50(0.46,0.55) | 0.48(0.43,0.54) |
| Sex | 0.02(0.02,0.02) | -0.01(-0.01,-0.01) | 0.05(0.01,0.08) | 0.08(0.05,0.12) | 0.00(-0.04,0.04) | 0.00(-0.04,0.05) |
| Education |  | 0.00(0.00,0.00) |  | 0.04(0.00,0.08) |  | -0.02(-0.05,0.02) |
| Smoking |  | 0.08(0.08,0.09) |  | 0.09(0.05,0.13) |  | 0.03(-0.01,0.08) |
| Exercise |  | -0.06(-0.07,-0.06) |  | -0.03(-0.07,0.01) |  | -0.12(-0.18,-0.06) |
| BMI |  | 0.25(0.24,0.25) |  | 0.17(0.13,0.21) |  | 0.06(-0.00,0.11) |

*Note:* Confidence Intervals (CI) including zero indicate a non-significant association; Coefficients are standardized; effects of a SD change in neuroticism scores on a SD change in FI scores; AO50 = The Australian Over 50's study; BMI = body mass index; FI = Frailty index; SATSA = The Swedish Adoption/Twin Study of Aging; UKB = UK Biobank; <sup>a</sup> - adjusted for age and sex, <sup>b</sup> - additionally adjusted for educational level, smoking status, exercise and BMI.

**Table 3.** The longitudinal association between baseline neuroticism and follow-up FI scores in SALT and SATSA cohorts (Beta and 95% CI).

|  | SALT |  | SATSA |  |  |  |
| --- | --- | --- | --- | --- | --- | --- |
|  | FI score 25 to 31-year follow-up |  | FI score 6-year follow-up |  | FI score 23-year follow-up |  |
|  | Model 1<sup>a</sup> | Model 2<sup>b</sup> | Model 1 | Model 2 | Model 1 | Model 2 |
|  | n = 18,960 | n = 18,773 | n = 1,031 | n = 1,031 | n = 418 | n = 418 |
| Neuroticism | 0.24(0.22,0.25) | 0.22(0.21,0.24) | 0.31(0.24,0.38) | 0.06(0.02,0.11) | 0.16(0.07,0.25) | 0.04(-0.05,0.12) |
| Age | 0.12(0.10,0.13) | 0.10(0.09,0.12) | 0.49(0.41,0.57) | 0.19(0.13,0.26) | 0.60(0.44,0.76) | 0.53(0.37,0.68) |
| Sex | 0.12(0.11,0.14) | 0.13(0.11,0.14) | 0.08(0.03,0.14) | 0.07(0.03,0.11) | 0.08(-0.01,0.16) | 0.08(0.01,0.16) |
| Education | -0.04(-0.06,-0.03) | -0.05(-0.06,-0.04) | 0.02(-0.03,0.06) | 0.02(-0.02,0.05) | -0.00(-0.07,0.06) | 0.00(-0.06,0.06) |
| Smoking | 0.03(0.02,0.04) | 0.03(0.01,0.04) | 0.04(-0.01,0.10) | 0.01(-0.03,0.06) | 0.08(0.00,0.16) | 0.07(0.00,0.14) |
| Exercise | -0.04(-0.05,-0.02) | -0.04(-0.05,-0.02) | -0.05(-0.12,0.01) | 0.00(-0.04,0.04) | 0.05(-0.02,0.13) | 0.02(-0.05,0.09) |
| BMI | 0.13(0.11,0.15) | 0.13(0.11,0.15) | 0.11(0.04,0.19) | 0.03(-0.02,0.08) | 0.08(-0.01,0.17) | 0.06(-0.02,0.14) |
| Baseline FI/<br>Chronic illness |  | 0.13(0.12,0.15) |  | 0.78(0.71,0.85) |  | 0.56(0.42,0.70) |

*Note:* Confidence Intervals (CI) including zero indicate a non-significant association; Coefficients are standardized; effects of a SD
change in neuroticism scores on a SD change in FI scores; BMI = body mass index; FI = Frailty index; SALT = Screening Across the
Lifespan of Twins Study; SATSA = The Swedish Adoption/Twin Study of Aging; <sup>a</sup> - adjusted for age, sex, educational level, smoking status, exercise and BMI <sup>b</sup> -additionally adjusted for baseline FI / Chronic illness.

Item-level sensitivity analysis showed that most neuroticism items were associated with frailty with no single item standing out across the four samples (*Supplementary Tables 5 and 6*). In addition, neuroticism sum-score was associated with most frailty items in all four samples, although relatively stronger associations were found for depressed mood and self-rated health (*Supplementary Tables 7 and 8*).

Within-pair differences in neuroticism significantly predicted within-pair differences in FI scores cross-sectionally, both in DZ and MZ twin pairs (Figure 1a). The association was lower in DZ pairs compared to the cross-sectional association observed in the full cohorts and even lower in MZ pairs, although the attenuation was not significant. Within-pair differences in baseline neuroticism also significantly predicted within-pair differences in follow-up FI scores. Again, there is a trend towards a weaker association between neuroticism and FI scores in MZ twins compared to DZ twins and compared to the association observed in the full cohort, especially evident in SALT (Figure 1b).

**Figure 1.**
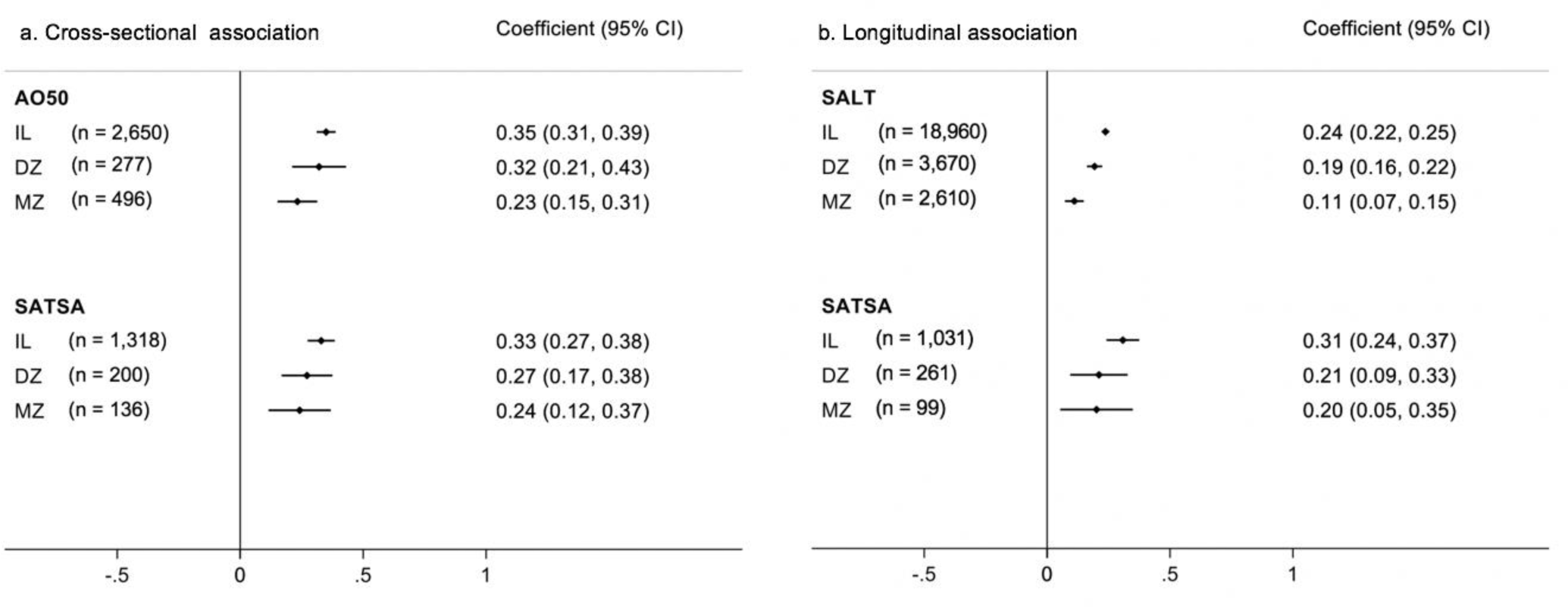
The cross-sectional (a) and longitudinal (b) associations between neuroticism and FI scores in AO50, SATSA and SALT for the individual level association observed in the full cohort as well as for same-sex DZ and MZ twins (Beta, 95% CI). *Note:* Models corrected for relatedness and covariates: education, smoking, exercise, and BMI; Longitudinal association in SALT over 25-29 years and in SATSA over six years; CI=Confidence interval; DZ=Dizygotic twins; IL=Individual level; MZ=Monozygotic twins; AO50=The Australian Over 50’s study; SATSA=The Swedish Adoption/Twin Study of Aging; SALT=Screening Across the Lifespan of Twins Study.

The best PRS_N_ explained 1.3% of the variance in neuroticism in UKB (*p*_T_ < 0.1), 0.5% in AO50 (*p*_T_ < 1×10^−5^), 0.3% in SALT (*p*_T_ < 0.3), and 1.8% in SATSA (*p*_T_ < 1) (*Supplementary Figure 2*). Further, PRS_N_ explain 0.36% of the FI variance in UKB and 0.26% in SALT but was non-significant in AO50 and SATSA (Figure 2). When meta-analyzed, overall higher polygenic risk for neuroticism significantly predicted FI scores (Figure 3). See *Supplementary Table 9* for sensitivity analysis including all eight *p*_T_s.

**Figure 2.**
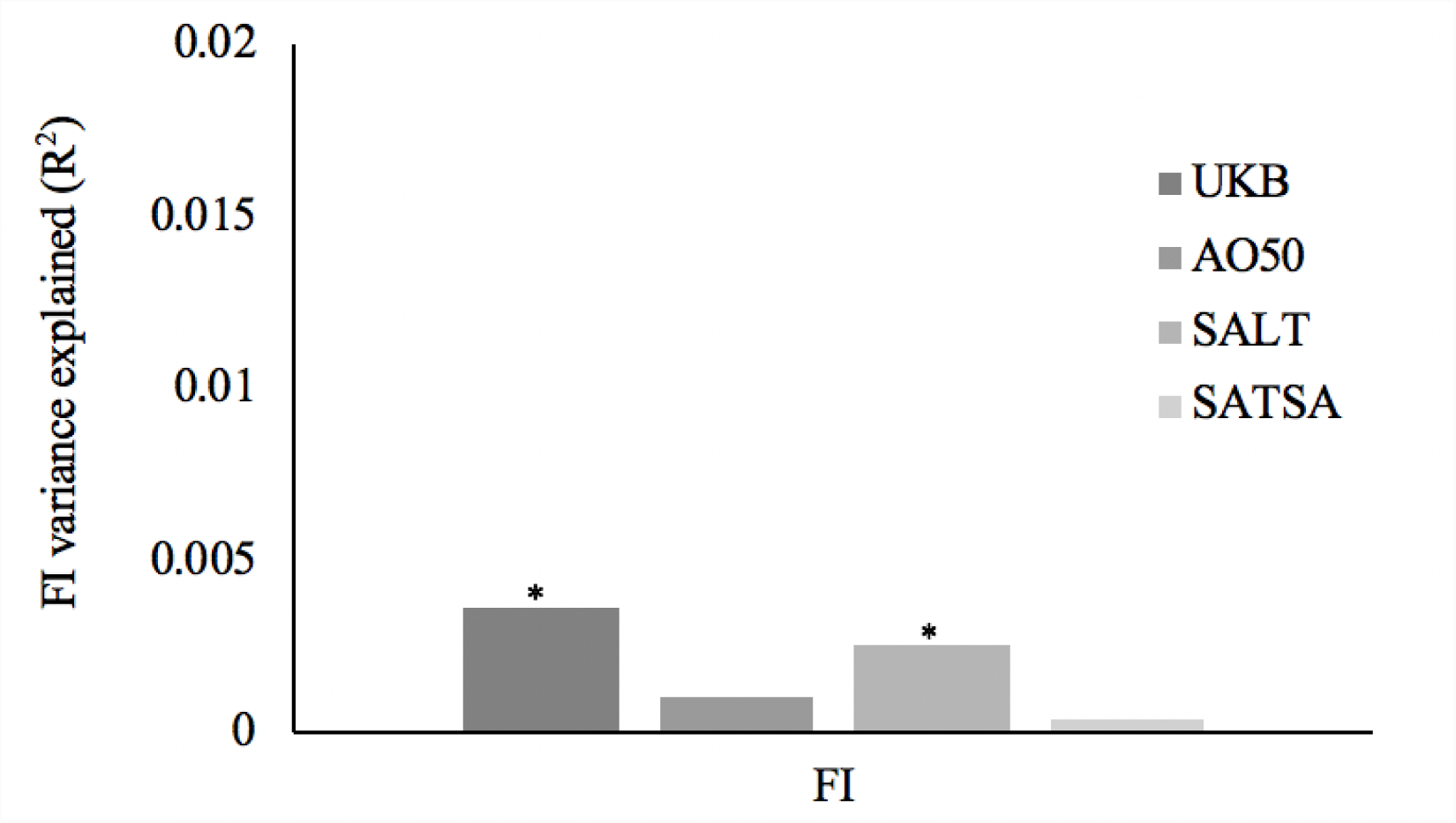
Variance in FI explained by polygenic risk scores for neuroticism in UKB (n = 243,734), AO50 (n =1,037), SALT (n = 6,221) and SATSA (n = 548) cohorts. *Note:* Variance refers to the difference in R^2^ between full and reduced regression models. AO50=The Australian Over 50’s study; CI=Confidence interval; FI=Frailty index; SALT=Screening Across the Lifespan of Twins Study; SATSA=The Swedish Adoption/Twin Study of Aging; UKB=UK Biobank; \**p*<0.001.

**Figure 3.**
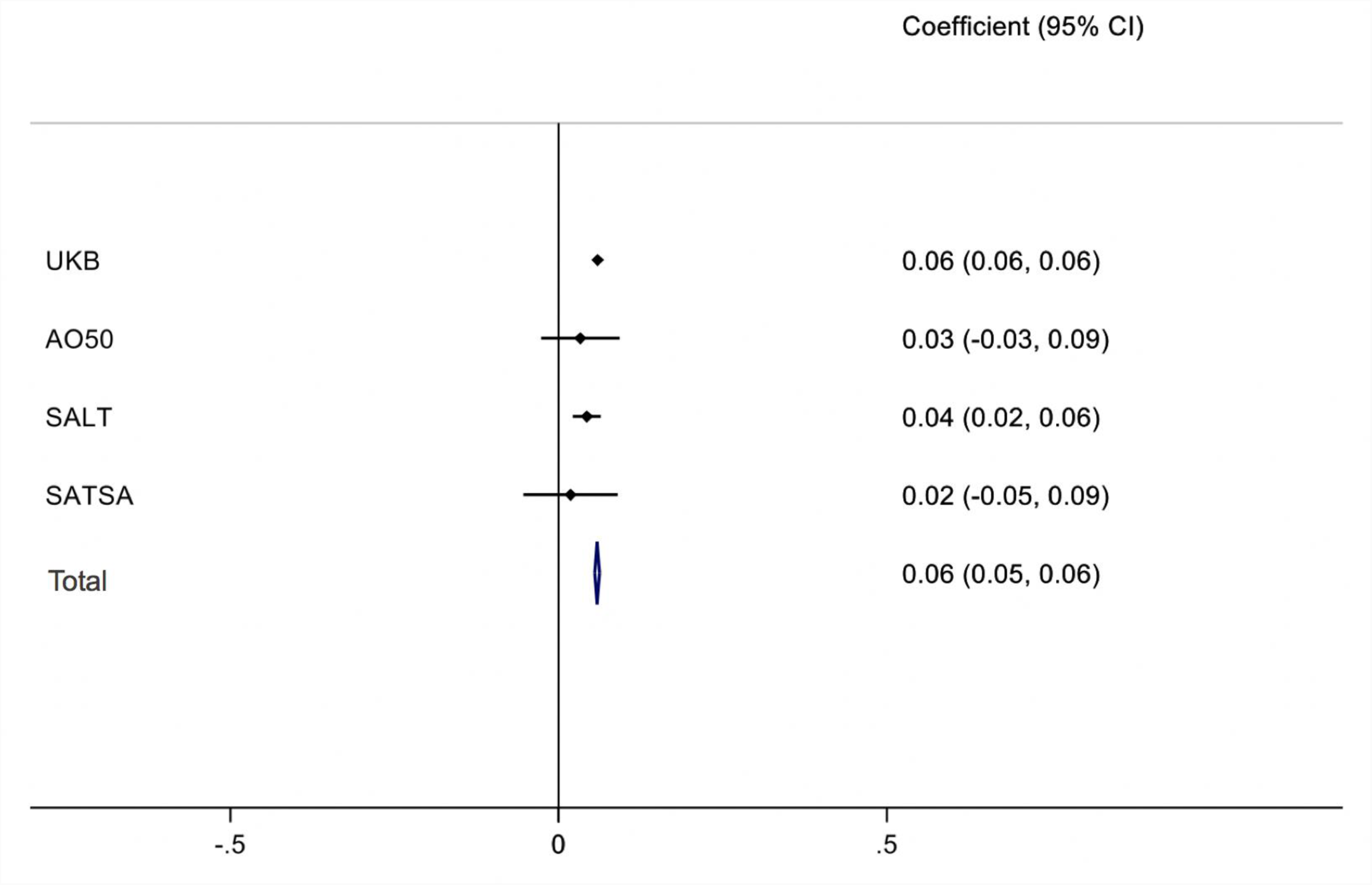
Effect sizes of polygenic risk scores for neuroticism on FI scores in UKB (n = 243,734), AO50 (n =1,037), SALT (n = 6,221) and SATSA (n = 548) cohorts and meta-analytical effect size combining the observed effect in the four cohorts and taking into account sample size (Beta and 95% CI). *Note*: Models were adjusted for age, sex, and PC’s; AO50=The Australian Over 50’s study; CI=Confidence interval; SALT=Screening Across the Lifespan of Twins Study; SATSA=The Swedish Adoption/Twin Study of Aging; UKB=UK Biobank.

## Discussion

By using cross-sectional and longitudinal data from large genetically informative samples of middle-aged and older adults, we found that higher neuroticism was consistently associated with greater frailty cross-sectionally and over more than two decades. After controlling for underlying genetic and shared environmental factors using a co-twin control design, the association between neuroticism and frailty remained evident, although attenuated to some extent, potentially indicating some shared underlying liability. Results from polygenic risk score analyses suggested the contribution of neuroticism-related genetic risk variants in frailty.

Overall, the results of the phenotypic analyses are in line with previous studies examining the association between neuroticism and frailty, using both measures of physical frailty (i.e. the frailty phenotype) and the frailty index (10-12). Our study expanded the follow-up time up to 29 years, elucidating the stability of neuroticism in midlife as a predictor of late life frailty. The associations between neuroticism and frailty were independent of age, sex, education and three lifestyle factors; smoking status, exercise and BMI, suggesting that neuroticism influences frailty over and above these potential confounding or mediating variables. To reduce the possibility of reverse causation by which poorer health at baseline may have influenced responses to items on the neuroticism scale, we additionally adjusted for baseline chronic illness/frailty. Although the effect size diminished, the association remained significant in SALT, and in SATSA across six years but not 23-years where attrition affected the power as indicated by the vastly reduced sample size.

Our second aim was to assess how familial influences contribute to the relationship between neuroticism and frailty. Using the co-twin control design, we found that the neuroticism-frailty association remained evident even when controlling for all underlying genetic and shared environmental factors (i.e. in MZ twins). This finding is in line with the notion that higher neuroticism increases the risk of frailty (29). However, comparison of effect size attenuation (though not significant) in DZ and even further in MZ twins also suggests that part of the association between neuroticism and frailty may be due to underlying shared factors, such as genetic risk.

Genetic risk for neuroticism was found to significantly predict frailty in the UKB and SALT, but not in AO50 and SATSA, which is likely due to the much smaller sample sizes and consequently low power. In all four cohorts, the amount of variance in FI scores explained by the PRS_N_ was small but not surprising considering the low predictive power of genetic risk scores in general (30). With increasing power of the discovery GWAS, estimation of effects sizes of common SNPs become more precise and PRS prediction will gain predictive power.

Together, our results demonstrate the involvement of both environmental and genetic factors in the relationship between neuroticism and health in late life. One possible mechanism through which neuroticism influences frailty is engagement in risky health behaviors. Previous research has shown that individuals with high neuroticism are more likely to smoke and have low physical activity (8, 31), both factors that have previously been associated with frailty (32, 33). Here, the neuroticism-frailty association only attenuated when adjusting for lifestyle factors. However, our measures were crude (binary) and there may be other unmeasured health-related behaviors that could influence frailty. Another possible explanation is that some mental health related aspects such as mood, feelings of loneliness or nervousness are reflected in both measures of neuroticism and frailty. However, results of the sensitivity analysis with re-calculated FI excluding all mental health items remained the same, deeming this explanation unlikely. Also, genetic overlap between neuroticism and frailty may contribute to the association and this should be investigated in the future when GWAS results on frailty become available. One possible biological mechanism through which neuroticism could potentially influence frailty is the Hypothalamic-Pituitary-Adrenal (HPA) axis activity. Physiological reserve is a prominent feature of frailty (2) and high neuroticism has been associated with dysregulation of the HPA axis (34). Future studies will need to examine the extent to which HPA dysregulation explains the neuroticism-frailty association.

This study has some limitations. First, FI was not available from SALT baseline measurement (Q73). The sample was relatively young at the time, and since prevalence of frailty is low in young people (2) there would be little variance in frailty. However, we used information about chronic or serious illness collected in 1973 to adjust for baseline health status. Second, an insufficient sample size may have been a limitation for some analyses, such as in the longitudinal analysis in SATSA with the largest time interval and the polygenic prediction in SATSA and AO50. However, with the use of several cohorts we could derive relatively consistent findings, highlighting the importance of well-powered samples, replication and meta-analytic methods, especially when using the PRS approach.

In conclusion, this study indicates that in addition to physical and biological determinants of frailty, psychological predictors of frailty should also be acknowledged. The results provide evidence that high neuroticism is associated with the development and course of frailty and that, although the association may in part reflect shared underlying genetic liability, neuroticism may significantly increase the risk for frailty.

## Ethical approvals

All participants have given informed consent. Ethical permits have been granted for UKB (16/NW/0274), AO50 (P1204), SALT (00-132), and SATSA (98-319) studies.

## Acknowledgments

We thank Xia Li for deriving the FI in SALT, Emma Raymond for her advice on FI methodology and Dr. Robert Karlsson for bioinformatics support with UKB data. We thank the Swedish and Australian twins for their participation and UKB participants. We also gratefully acknowledge the support and funding for the Over-50 twin study by Mr. George Landers of Chania, Crete. We acknowledge The Swedish Twin Registry for access to data. The Swedish Twin Registry is managed by Karolinska Institutet and receives funding through the Swedish Research Council under the grant no 2017-00641. In addition, UKB data was used under application number 22224.

