## Supplementary Material for "Neuroticism as a predictor of frailty in old age: a genetically informative approach"

### Supplementary Materials

#### Supplementary Method

*Frailty Index in AO50 study*

Supplementary Table 1 lists the 41 health deficits selected for the FI in the AO50. Information from participants with less than 20% missing answers was included in the calculation of the FI and all items included in the index had less than 10% missing data. Of the total 41 items, 16 items had missing data points and the pattern of missingness for these variables was examined. After that, multiple imputation was used to replace the missing values in order to maximize the utilization of the data. We performed 56 rounds of imputation and the pooled mean from the simulations was used as the final value for each missing data point. Next, the FI was calculated by adding up the number of deficits for each individual and dividing the sum by the 41 items included (e.g. FI = 14/41 = 0.34) resulting in a score ranging from 0 to 1, where higher values indicate greater frailty. It should be noted that although the theoretical maximum of the FI is 1, >99% of individuals in all populations have an FI< 0.7, indicating that survival beyond this point is lethal (1). To validate the FI in AO50, we assessed associations with age, sex, and risk of all-cause mortality (follow-up ≤ 19 years). Women had on average higher FI scores than men and there was a curvilinear trend toward higher values in older participants (data not shown), consistent with previously reported FI´s (2-5). In addition, we performed a Cox regression analysis for all-cause mortality using the sum of the FI item scores across the 41 items as the independent variable (so that the HR is interpretable as the risk associated with increase in one deficit) and age, BMI, and smoking status as covariates and stratifying by sex. The FI was independently associated with increased risk for all-cause mortality in both women (HR per accumulation of one deficit 1.05, 95% CI 1.02-1.08), and men (HR 1.07, 95% CI 1.04-1.11). We also performed a sensitivity analysis and examined the association between FI and all-cause mortality for those participants with no missing data in the FI items (n = 2275) and results were almost identical with those from the imputed dataset (data not shown).

Supplementary Table 1. The 41 items included in the FI in AO50

| No. | Frailty index item | Coding |
|---|---|---|
| 1 | Hearing problems | No=0, Yes=1 |
| 2 | Glaucoma | No=0, Yes=1 |
| 3 | Recent health decline and mobility loss | No=0, Yes=1 |
| 4 | Self-rated health | Very good=0, Fair=0.5, Poor/Very poor=1 |
| 5 | Cancer / Leukaemia | No=0, Yes=1 |
| 6 | Chronic bronchitis or emphysema | No=0, Yes=1 |
| 7 | Cataracts | No=0, Yes=1 |
| 8 | Chest pain / Angina | No=0, Yes=1 |
| 9 | Diabetes | No=0, Yes=1 |
| 10 | Thyroid disease | No=0, Yes=1 |
| 11 | Heart attack | No=0, Yes=1 |
| 12 | High blood pressure | No=0, Yes=1 |
| 13 | Kidney disease | No=0, Yes=1 |
| 14 | Osteoporosis | No=0, Yes=1 |
| 15 | Anemia | No=0, Yes=1 |
| 16 | Stroke | No=0, Yes=1 |
| 17 | Stomach ulcer | No=0, Yes=1 |
| 18 | Eczema | No=0, Yes=1 |
| 19 | Asthma | No=0, Yes=1 |
| 20 | Able to perform self-care | Completely able=0, Some limitations/Very limited |
| 21 | Able to walk | Completely able=0, Some limitations=0.5, Very limited |
| 22 | Incontinence | No=0, Yes=1 |
| 23 | Able to perform work and housekeeping | Completely able=0, Some limitations=0.5, Very limited |
| 24 | Able to perform recreational and leisure time activities | Completely able=0, Some limitations=0.5, Very limited |
| 25 | Loneliness | No=0, Yes=1 |
| 26 | Unhappy and depressed | Not at all=0, No more than usual=0.5, Rather more than usual/much more than usual=1 |
| 27 | Happiness | More so than usual=0, Same as usual=0.5, Less than usual/Much less than usual=1 |
| 28 | Feeling tired after physical activity | No=0, Yes=1 |
| 29 | Muscles tired after physical activity | No=0, Yes=1 |
| 30 | Migraine | No=0, Yes=1 |
| 31 | Joint pain | No=0, Yes=1 |
| 32 | Current back pain | No=0, Yes=1 |

Supplementary Table 1 (continued)

| | | |
|---|---|---|
| 33 | Current hip pain | No=0, Yes=1 |
| 34 | Current knee pain | No=0, Yes=1 |
| 35 | Recurrent abdominal pain | No=0, Yes=1 |
| 36 | Gallbladder trouble | No=0, Yes=1 |
| 37 | High cholesterol | No=0, Yes=1 |
| 38 | Gout | No=0, Yes=1 |
| 39 | Hernia or rupture | No=0, Yes=1 |
| 40 | Liver disease | No=0, Yes=1 |
| 41 | Fatigue | 0=Not at all, 0.5=A little, 1=A lot/ Unbearably |

Supplementary Figure 1. Distribution of neuroticism scores in UKB (n = 401.655), AO50 (n = 2.946), SALT (n = 21.175), and SATSA (n = 1.575) cohorts.

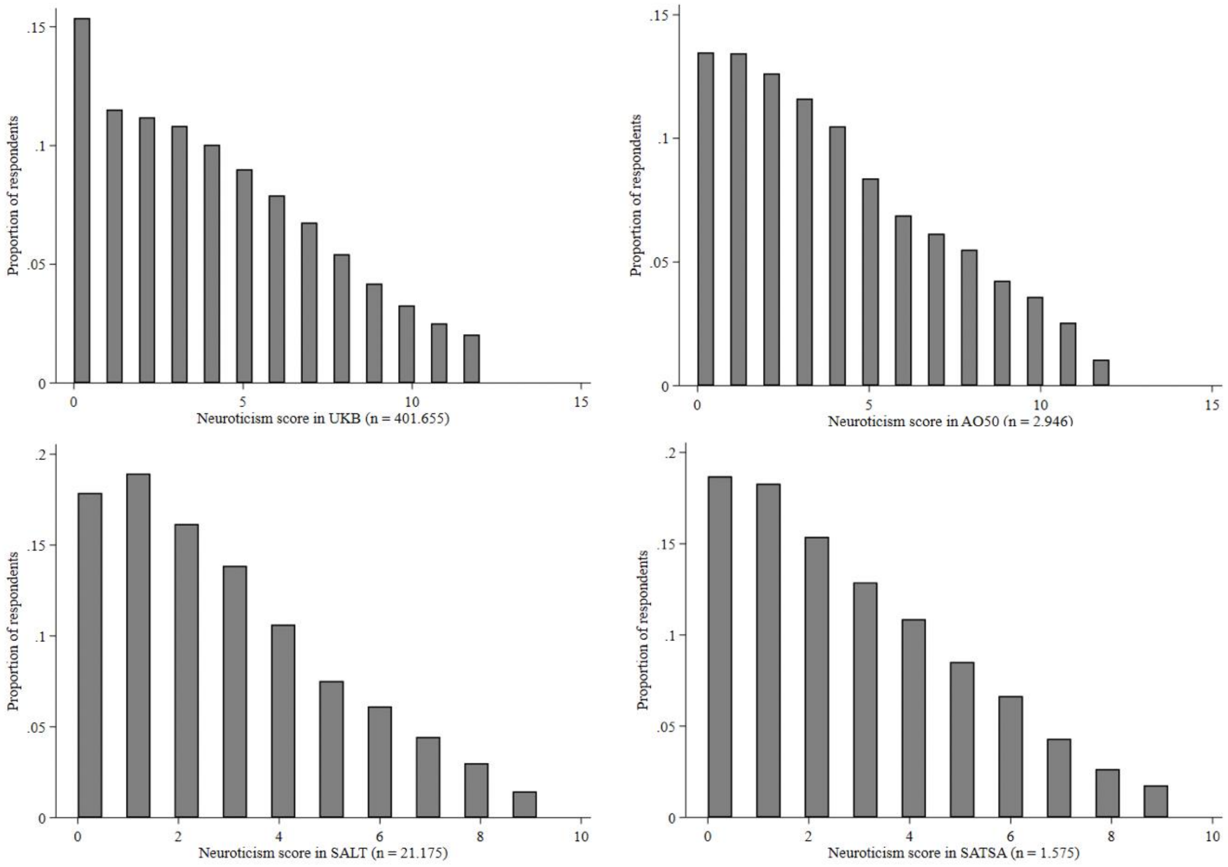

Supplementary Table 2. Items excluded from FI´s due to overlap with the respective EPQ scales

| Overlapping item | Coding |
|---|---|
| **UKB** | |
| Miserableness | No=0, Yes=1 |
| Loneliness | No=0, Yes=1 |
| Nervous personality | No=0, Yes=1 |
| **AO50** | |
| Loneliness | No=0, Yes=1 |
| **SATSA** | |
| Usually feels tired | No=0, Yes=1 |
| Consider oneself happy and carefree | No=1, Yes=0 |

Supplementary Table 3. Items excluded from additional FI´s for sensitivity analysis without any mental health variables

| Mental health item | Coding |
|---|---|
| **UKB** | |
| Severe anxiety | No=0, Yes=1 |
| Depressed mood | Not at all=0, Several days=0.25, More than half=0.5, Nearly every day=1 |
| Tiredness/Lethargy | Not at all=0, Several days=0.25, More than half=0.5, Nearly every day=1 |
| Sleeplessness/Insomnia | Never=0, Rarely=0.5, Sometimes/Usually=1 |
| **AO50** | |
| Unhappy and depressed | Not at all=0, No more than usual=0.5, Rather more than usual/much more than usual=1 |
| Happiness | Better than usual=0, Same as usual=0.5 Less than usual/Much less than usual=1 |
| Fatigue | More so than usual=0, Same as usual=0.5, Less than usual/Much less than usual=1 |
| **SATSA** | |
| Feeling lonely | Never, Almost never, Rather seldom=0, Quite often, Always, Almost always=1 |
| Feeling depressed | Never, Almost never, Rather seldom=0, Quite often, Always, Almost always=1 |

*Computing polygenic risk scores for neuroticism*

For all four samples, PRS were calculated based on the same summary statistics from GWAS meta-analyses (6). Sample overlap between the discovery and the target sample may possibly lead to inflation of PRS effects and therefore sample overlap should be checked and addressed if needed. In this study, the sample overlap between discovery and target samples were dealt with by using the leave-one-out summary statistics (AO50 and SALT) or by excluding overlapping individuals and their relateds (UKB and SATSA).

Since the discovery sample included the UKB interim release genetic data with up to 107,245 individuals, we removed all individuals in the interim release and individuals related to participants in the interim release up to third degree from the target sample (total n removed with this step = 191,092). In addition to the initial quality control conducted centrally, further variants were excluded based on strand ambiguity, low imputation quality (INFO < 0.8) and being triallelic SNPs. Samples were excluded based on recommended exclusions (poor heterozygosity / missingness), if not being in the white British ancestry subset and if participation consent had been withdrawn. Clumping was used to select the most significantly associated SNPs and exclude those in strong linkage disequilibrium (LD) (--clump flag in Plink 2.0 with $r^2$ < 0.1 within a 1000 kb window). PRS were computed under eight *p*-value thresholds, ranging from $5e^{-8}$ to 1 (--score command in Plink 2.0) and standardized for further analysis.

In AO50 study, the SNPs with low imputation quality (r2<0.6) and MAF below 1% were excluded and most significant independent SNPs were selected using PLINK1.9 (clumping criteria LD r2<0.1 within windows of 10 mb). Eight different PRS were calculated using different *p*-value thresholding of the GWAS summary statistics, ranging from $5e^{-8}$ to 1.

In SALT, the scores were derived in imputed dosage data after filtering out SNPs with MAF<0.05 and INFO<0.8. LD-clumping was run using the 1000-Genomes reference population, to obtain a relatively independent set of autosomal SNPs for each set of discovery results, while retaining the most significant SNP in each LD block and maximizing the overlap of SNPs across the discovery and target data. The following parameters were applied in PLINK.v.1.9: --clump-kb 1000 --clump-r2 0.1, i.e., removing any SNPs that are in $r2 \geq 0.1$ and within 1MB window from the index SNP. PRS were calculated under eight *p*-value thresholds, ranging from 0.001 to 1.

Detailed information of the genotyping and creation of the $PRS_N$ in SATSA can be found elsewhere (7). The maximum number of LD-pruned SNPs considered in computing $PRS_N$ in each sample were as follows: 206 340 in UKB; 239 352 in AO50; 110 678 in SALT, 144 367 in SATSA.

*Covariate assessment*

In SALT, education measured at follow-up was used as a covariate, since participants were fairly young (youngest were 15 years old) at the baseline measurement and many had not completed their final educational degree at that time point. Education was reported in years in UKB and AO50. In UKB, education was coded as a dichotomous variable indicating the educational attainment level above the compulsory (0 – compulsory; 1 – above compulsory). In SATSA, education was measured on a scale from 1 (elementary school) to 4 (university level or higher) and in SALT, education was assessed with a two-point scale, 0 (compulsory education) and 1 (more than compulsory education achieved). To be consistent across cohorts, educational level was dichotomized into low versus high education. In UKB, exercise was measured with two items assessing the frequency of moderate and vigorous exercise a week (ranging from 0 to 7 times a week). In AO50, exercise was assessed on a scale from 1 (nil) to 4 (high) and in SATSA on a scale ranging from 1 (I hardly get any exercise at all) to 7 (I get very much exercise). Again, in order to be consistent across cohorts, exercise was dichotomized into exercisers and non-exercisers. In all cohorts, smoking status was classified into the following three groups: non-smokers, ex-smokers, and current smokers.

*Statistical analysis*

To avoid excluding observations because of genetic relatedness in UK Biobank and to retain maximum power, a family ID was created based on available genetic relatedness information. To obtain a family ID in UK Biobank, the KING algorithm (8) was used to identify pairs of genetically related individuals using 0.0884 as the kinship cut-off (second degree relatives and closer). No closely related individuals were found for 419,467 participants and 11 closely related individuals were identified on one occasion (Cluster sizes: 1= 419467; 2= 27654; 3= 3442; 4= 586; 5= 125; 6= 32; 7= 11; 8= 1; 9= 1;10= 1; 11= 1). Each genetically related cluster was assigned a Family ID code and was used in all analyses for adjusting for relatedness between participants by applying the ‘sandwich’ estimator.

Supplementary Table 4. The phenotypic cross-sectional association between neuroticism and FI scores without mental health items in UKB, AO50 and SATSA cohorts (Beta and 95% CI)

| | UKB Coefficient (95% CI) | AO50 Coefficient (95% CI) | SATSA Coefficient (95% CI) |
|---|---|---|---|
| | n = 323.136 | n = 2.650 | n = 1.318 |
| Neuroticism | 0.242 (0.239, 0.246) *** | 0.293 (0.254, 0.332) *** | 0.289 (0.231, 0.347) *** |
| Age | 0.186 (0.183, 0.189) *** | 0.284 (0.242, 0.326) *** | 0.470 (0.413, 0.527) *** |
| Sex | 0.005 (0.002, 0.008) ** | 0.089 (0.053, 0.125) *** | -0.011 (-0.057, 0.034) |
| Education | 0.004 (0.001, 0.007) * | 0.040 (0.002, 0.079) * | -0.020 (-0.052, 0.013) |
| Smoking Status | 0.081 (0.078, 0.085) *** | 0.088 (0.051, 0.125) *** | 0.027 (-0.017, 0.072) |
| Exercise | -0.057 (-0.061, -0.054) *** | -0.032 (-0.071, 0.007) | -0.128 (-0.189, -0.067) *** |
| BMI | 0.249 (0.245, 0.252) *** | 0.175 (0.135, 0.216) *** | 0.053 (-0.005, 0.110) |

Coefficients are standardized; effects of a SD change in neuroticism scores on a SD change in FI scores

*$p$<0.05; **$p$<0.01; ***$p$<0.001

Supplementary Table 5. The cross-sectional association between individual neuroticism item and FI score in UKB, AO50 and SATSA cohorts (Beta and 95% CI)

| | UKB Coefficient (95% CI) | AO50 Coefficient (95% CI) | SATSA Coefficient (95% CI) |
|---|---|---|---|
| | n = 395.838 | n = 2.438 | n = 1.258 |
| **Neuroticism Item** | | | |
| Mood up and down | 0.086 (0.082, 0.090) *** | 0.071 (0.025, 0.116) ** | 0.033 (-0.020, 0.086) |
| Feeling miserable | 0.073 (0.069, 0.076) *** | 0.030 (-0.012, 0.072) | na |
| Irritable person | 0.011 (0.007, 0.014) *** | 0.068 (0.027, 0.110) ** | na |
| Easily hurt | 0.010 (0.007, 0.013) *** | 0.011 (-0.027, 0.049) | 0.061 (0.011, 0.110) * |
| Feel fed-up | 0.077 (0.074, 0.081) *** | 0.056 (0.012, 0.100) * | na |
| Nervous person | 0.020 (0.017, 0.024) *** | 0.023 (-0.030, 0.075) | na |
| Anxious / Worrier | 0.052 (0.049, 0.056) *** | 0.074 (0.031, 0.117) ** | 0.063 (0.001, 0.124) * |
| Tense | 0.044 (0.041, 0.048) *** | 0.095 (0.045, 0.146) *** | na |
| Suffer nerves | 0.033 (0.030, 0.036) *** | 0.050 (0.002, 0.099) * | 0.152 (0.049, 0.255) ** |
| Feel lonely | 0.070 (0.066, 0.073) *** | 0.049 (0.006, 0.093) * | na |
| Worry long after embarrassment | 0.011 (0.008, 0.014) *** | 0.040 (-0.002, 0.083) | -0.043 (-0.091, 0.005) |
| Guilty feelings | 0.030 (0.026, 0.033) *** | 0.025 (-0.015, 0.065) | na |
| Often make decisions late | na | na | 0.038 (-0.019, 0.094) |
| Feel tired / ill for no reason | na | na | 0.171 (0.105, 0.238) *** |
| Often deep in thought | na | na | 0.056 (0.007, 0.106) * |
| Very restless | na | na | 0.004 (-0.044, 0.052 |
| **Covariates** | | | |
| Age | 0.176 (0.173, 0.178) *** | 0.263 (0.220, 0.306) *** | 0.447 (0.389, 0.505) *** |
| Sex | -0.021 (-0.024, -0.018) *** | 0.093 (0.057, 0.130) *** | 0.027 (-0.018, 0.073) |
| Education | 0.001 (-0.001, 0.004) | 0.037 (-0.003, 0.077) | -0.008 (-0.041, 0.024) |
| Exercise | -0.070 (-0.074, -0.068) *** | -0.034 (-0.075, 0.007) | -0.127 (-0.188, -0.066) *** |
| Smoking | 0.081 (0.078, 0.083) *** | 0.089 (0.052, 0.126) *** | 0.026 (-0.019, 0.070) |
| BMI | 0.238 (0.235, 0.241) *** | 0.171 (0.129, 0.213) | 0.052 (-0.008, 0.111) |

Coefficients are standardized; effects of a SD change in individual neuroticism item score on a SD change in FI scores
*$p$<0.05; **$p$<0.01; ***$p$<0.001

Supplementary Table 6. The longitudinal association between baseline individual neuroticism item and follow-up FI score (Beta and 95% CI)

| | SALT<br>Coefficient (95% CI) |
|---|---|
| | n = 18.157 |
| **Neuroticism Item** | |
| Mood up and down | 0.011 (-0.003, 0.026) |
| Extra sensitive | 0.049 (0.034, 0.064) *** |
| Anxious / Worrier | 0.038 (0.021, 0.055) *** |
| Suffer nerves | 0.107 (0.089, 0.125) *** |
| Worry long after embarrassment | 0.027 (0.012, 0.042) ** |
| Often make decisions late | 0.003 (-0.011, 0.017) |
| Feel tired / ill for no reason | 0.076 (0.060, 0.092) *** |
| Often deep in thought | 0.028 (0.013, 0.043) *** |
| Very restless | 0.034 (0.019, 0.049) *** |
| **Covariates** | |
| Age | 0.097 (0.080, 0.114) *** |
| Sex | 0.130 (0.115, 0.146) *** |
| Education | -0.045 (-0.058, -0.033) *** |
| Exercise | -0.035 (-0.050, -0.020) *** |
| Smoking | 0.027 (0.014, 0.040) *** |
| BMI | 0.128 (0.109, 0.146) *** |
| Baseline FI /Chronic illness | 0.129 (0.114, 0.144) *** |

Coefficients are standardized; effects of a SD change in individual neuroticism item score on a SD change in FI scores
*$p$<0.05; **$p$<0.01; ***$p$<0.001

Supplementary Table 7. The cross-sectional association between neuroticism and individual FI items in UKB, AO50 and SATSA cohorts (Beta and 95% CI)

| FI item | UKB Coefficient (95% CI) | AO50 Coefficient (95% CI) | SATSA Coefficient (95% CI) |
|---|---|---|---|
| | n = 314.132 to 323.136 | n = 2.650 | n = 1.318 |
| Glaucoma | 0.008 (0.005, 0.011) *** | 0.017 (-0.027, 0.061) | |
| Cataracts | 0.017 (0.014, 0.021) *** | 0.021 (-0.017, 0.059) | 0.029 (-0.030, 0.087) |
| Hearing problems | 0.098 (0.095, 0.102) *** | 0.059 (0.021, 0.096) ** | 0.092 (0.038, 0.146) ** |
| Vision problems | | | 0.156 (0.099, 0.214) *** |
| Migraine | 0.042 (0.038, 0.045) *** | 0.123 (0.081, 0.164) *** | |
| Dental problems | 0.117 (0.113, 0.120) | | |
| Recent health decline and mobility loss | | 0.158 (0.114, 0.203) *** | |
| Health prevents from doing things normally would like to do | | | 0.258 (0.201, 0.315) *** |
| Self-rated health | 0.231 (0.228, 0.235) *** | 0.198 (0.155, 0.241) *** | 0.298 (0.241, 0.355) *** |
| Able to perform self-care | | 0.075 (0.037, 0.112) *** | 0.082 (-0.00, 0.163) |
| Shower and bathe[a] | | | 0.058 (-0.001, 0.117) |
| Get in and out of bed[a] | | | 0.045 (-0.027, 0.117) |
| Dress and undress[a] | | | 0.036 (-0.030, 0.103) |
| Able to walk | | 0.089 (0.050, 0.128) *** | 0.056 (-0.002, 0.114) |
| Able to perform work and housekeeping | | 0.104 (0.063, 0.144) *** | 0.089 (0.035, 0.144) ** |
| Grocery shopping | | | 0.089 (0.032, 0.146) ** |
| Able to perform recreational and leisure time activities | | 0.126 (0.086, 0.166) *** | |
| Travel further distances[b] | | | 0.101 (0.046, 0.156) *** |
| Prepare meals[b] | | | 0.105 (0.048, 0.161) *** |
| Manage medications[b] | | | 0.093 (0.030, 0.157) ** |
| Manage money[b] | | | 0.109 (0.045, 0.173) ** |
| Use telephone[b] | | | 0.116 (0.058, 0.174) *** |
| Depressed | 0.498 (0.494, 0.501) *** | 0.457 (0.422, 0.492) *** | 0.542 (0.487, 0.598) *** |
| Unhappiness | | 0.159 (0.116, 0.203) *** | |
| Severe anxiety | 0.112 (0.108, 0.115) *** | | |

Supplementary Table 7 (continued.)

| | | | |
|---|---|---|---|
| Loneliness | | | 0.099 (0.040, 0.158) ** |
| Sleep / Insomnia | 0.261 (0.257, 0.264) *** | | |
| Fatigue | 0.340 (0.337, 0.344) *** | 0.331 (0.282, 0.380) *** | |
| Dizzyness | | | 0.225 (0.164, 0.287) *** |
| Infirmity | 0.130 (0.127, 0.133) *** | | |
| Falls | 0.091 (0.088, 0.095) *** | | |
| Fractures | 0.017 (0.013, 0.020) *** | | |
| Diabetes | 0.016 (0.013, 0.020) *** | 0.034 (-0.004, 0.072) | 0.098 (0.035, 0.161) ** |
| Heart attack | 0.015 (0.012, 0.018) *** | 0.080 (0.041, 0.119) *** | 0.060 (-0.005, 0.126) |
| Chest pain / Angina | 0.034 (0.030, 0.037) *** | 0.087 (0.047, 0.126) *** | 0.070 (0.012, 0.128) * |
| Chest pain | 0.143 (0.139, 0.146) *** | | |
| Stroke | 0.015 (0.012, 0.018) *** | 0.054 (0.013, 0.095) * | 0.076 (0.013, 0.140) * |
| High blood pressure | 0.060 (0.056, 0.063) *** | 0.100 (0.060, 0.140) *** | 0.060 (0.002, 0.119) * |
| Hypothyroidism | 0.027 (0.024, 0.030) *** | | |
| Deep-vein thrombosis | 0.006 (0.003, 0.009) *** | | |
| Circulation problems in arms/legs | | | 0.194 (0.129, 0.260) *** |
| High cholesterol | 0.030 (0.026, 0.033) *** | 0.077 (0.035, 0.119) *** | |
| Wheeze | 0.098 (0.095, 0.102) *** | | |
| Persistent cough | | | 0.091 (0.029, 0.154) ** |
| Pneumonia | 0.006 (0.003, 0.010) ** | | |
| Chronic bronchitis or emphysema | 0.032 (0.028, 0.035) *** | 0.081 (0.037, 0.125) *** | 0.094 (0.027, 0.162) ** |
| Asthma | 0.042 (0.038, 0.045) *** | 0.046 (0.005, 0.087) * | 0.036 (-0.024, 0.096) |
| Rheumatoid arthritis | 0.005 (0.002, 0.008) *** | | 0.068 (0.003, 0.132) * |
| Osteoarthritis | 0.034 (0.031, 0.038) *** | | 0.081 (0.015, 0.146) * |
| Gout | 0.002 (-0.001, 0.005) | 0.015 (-0.023, 0.053) | 0.042 (-0.019, 0.104) |
| Goiter and other gland problems | | | 0.042 (-0.019, 0.104) |
| Osteoporosis | 0.017 (0.014, 0.021) *** | 0.038 (-0.004, 0.080) | 0.068 (-0.005, 0.142) |
| Eczema / allergic rhinitis /allergic manifestations | 0.053 (0.050, 0.057) *** | 0.096 (0.055, 0.138) *** | 0.101 (0.042, 0.160) ** |
| Psoriasis | 0.015 (0.011, 0.018) *** | | |
| Any cancer or leukemia diagnosed | 0.001 (-0.003, 0.004) | 0.038 (-0.002, 0.078) | 0.009 (-0.051, 0.070) |

Supplementary Table 7 (continued.)

| | | | |
|---|---|---|---|
| Multiple cancer diagnosed | -0.002 (-0.005, 0.001) | | |
| Feeling tired after physical activity | | 0.258 (0.219, 0.297) *** | |
| Muscles tired after physical activity | | 0.275 (0.238, 0.312) *** | |
| Joint pain | | 0.142 (0.104, 0.181) *** | |
| Back pain | 0.126 (0.122, 0.129) *** | 0.065 (0.026, 0.105) ** | |
| Hip pain | 0.079 (0.075, 0.082) *** | 0.068 (0.025, 0.110) ** | |
| Knee pain | 0.087 (0.084, 0.090) *** | 0.098 (0.059, 0.138) *** | |
| Head and/or neck pain | 0.178 (0.174, 0.181) *** | | |
| Stomach/abdominal pain | 0.135 (0.131, 0.138) *** | 0.094 (0.046, 0.142) *** | |
| Stomach ulcer | | 0.068 (0.025, 0.111) ** | |
| Gastric ulcer | | | 0.083 (0.022, 0.143) ** |
| Whole body pain | 0.044 (0.040, 0.047) *** | | |
| Facial pain | 0.065 (0.061, 0.069) *** | | |
| Sciatica | 0.015 (0.011, 0.018) *** | | 0.144 (0.080, 0.209) *** |
| Hernia | 0.032 (0.028, 0.035) *** | 0.069 (0.029, 0.109) ** | |
| Incontinence | | 0.058 (0.018, 0.098) ** | 0.171 (0.099, 0.243) *** |
| Kidney disease | | 0.029 (-0.009, 0.067) | 0.045 (-0.016, 0.105) |
| Liver disease | | 0.014 (-0.027, 0.056) | |
| Thyroid disease | | 0.031 (-0.011, 0.072) | |
| Gastric reflux | 0.051 (0.048, 0.055) *** | | |
| Diverticulosis | 0.021 (0.018, 0.024) *** | | |
| Gall stones | 0.010 (0.007, 0.014) *** | | |
| Anemia | | 0.063 (0.021, 0.105) ** | 0.052 (-0.009, 0.113) |
| Gallbladder trouble | | 0.025 (-0.015, 0.065) | |

Coefficients are standardized; effects of a SD change neuroticism scores on a SD change in individual FI items scores

*$p$<0.05; **$p$<0.01; ***$p$<0.001

Supplementary Table 8. The longitudinal association between neuroticism (total score) and individual FI items in SALT (Beta and 95% CI)

| | SALT<br>Coefficient (95% CI) |
|---|---|
| | n = 18.773 |
| **FI item** | |
| General health (SRH) | 0.163 (0.148, 0.178) *** |
| Health prevent action | 0.129 (0.114, 0.145) *** |
| Infections (how often) | 0.055 (0.037, 0.073) *** |
| Buzzing in ears | 0.073 (0.057, 0.089) *** |
| Angina | 0.059 (0.045, 0.074) *** |
| Heart attack | 0.018 (0.005, 0.032) * |
| Heart failure | 0.019 (0.005, 0.033) * |
| High blood pressure | 0.028 (0.013, 0.044) *** |
| Lipid disorder (e.g. high cholesterol) | 0.036 (0.020, 0.052) *** |
| Vascular spasms in legs | 0.047 (0.032, 0.061) *** |
| Clot in the leg (Venus thrombosis) | 0.019 (0.005, 0.034) *** |
| Stroke | 0.019 (0.005, 0.032) *** |
| TIA attacks | 0.016 (0.001, 0.031) * |
| Do you have, or have you had irregular cardiac rhythm/atrial fibrillation | 0.057 (0.041, 0.072) *** |
| Bronchitis / emphysema | 0.068 (0.052, 0.084) *** |
| Dizziness | 0.140 (0.125, 0.155) *** |
| Rheumatoid arthritis | 0.021 (0.007, 0.037) * |
| Knee joint problem | 0.055 (0.041, 0.071) *** |
| Sciatica | 0.089 (0.073, 0.105) *** |
| Osteoporosis | 0.017 (0.002, 0.032) * |
| Hip joint problem | 0.058 (0.043, 0.074) *** |
| Back pain | 0.086 (0.070, 0.101) *** |
| Neck pain | 0.129 (0.113, 0.145) *** |
| Diabetes | 0.019 (0.004, 0.037) * |
| Goiter | 0.018 (0.003, 0.033) * |
| Glandular diseases (excluding goiter) | 0.018 (0.001, 0.034) * |
| Gall bladder problem | 0.033 (0.018, 0.048) *** |

Supplementary Table 8 (continued)

| | |
|---|---|
| Liver disease | 0.020 (0.004, 0.036) * |
| Gout | 0.023 (0.008, 0.039) * |
| Kidney disease | 0.016 (0.001, 0.031) * |
| Stomach problems | 0.135 (0.119, 0.150) *** |
| Recurring urinary tract problems | 0.083 (0.068, 0.098) *** |
| Cancer | 0.001 (-0.014, 0.015) |
| Migraine | 0.066 (0.050, 0.082) *** |
| Asthma | 0.049 (0.033, 0.066) *** |
| Allergies | 0.077 (0.061, 0.093) *** |
| Recurrent periods of coughing | 0.074 (0.057, 0.091) *** |
| Depressed | 0.211 (0.194, 0.229) *** |
| Happiness | 0.139 (0.121, 0.156) *** |
| Loneliness | 0.152 (0.136, 0.169) *** |
| Any physical handicaps | 0.046 (0.030, 0.062) *** |
| Crohn's disease or Ulcerative colitis | 0.037 (0.019, 0.055) *** |
| Eyesight | 0.062 (0.046, 0.077) *** |
| Hearing | 0.056 (0.041, 0.070) *** |

Coefficients are standardized; effects of a SD change neuroticism scores on a SD change in individual FI items scores

*$p<0.05$; **$p<0.01$; ***$p<0.001$

Supplementary Figure 2. PRS for neuroticism explaining variance in phenotypic neuroticism in UKB (n = 198.078), AO50 (n = 1.022), SALT (n = 5.832) and SATSA (n= 573) cohorts.

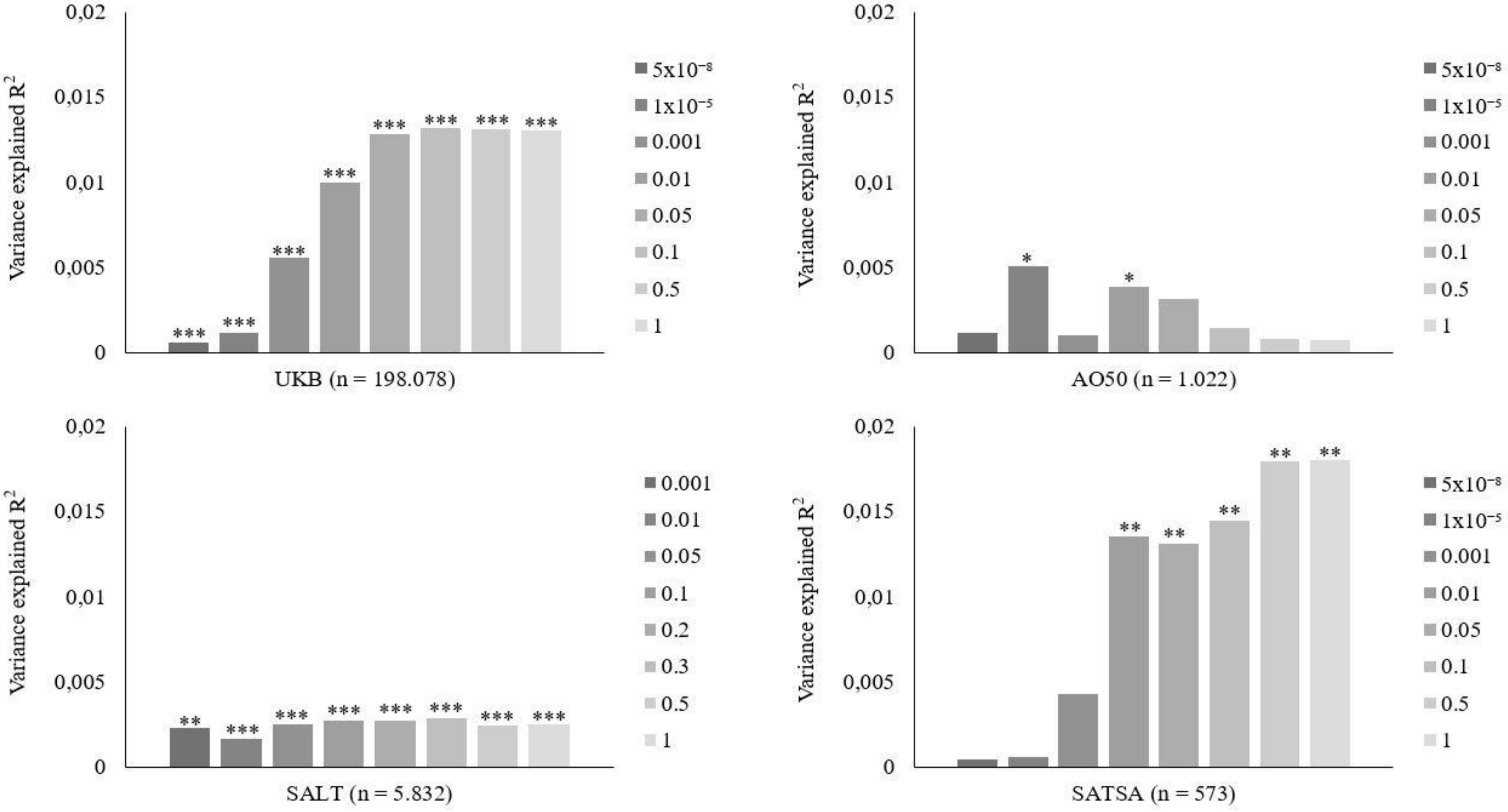


Note: *$p$<0.05; **$p$<0.01; ***$p$<0.001

Supplementary Table 9. Effects of $PRS_N$ on FI scores while adjusting for age, sex and PC´s (Beta and 95% CI)

| | UKB | AO50 | SALT | SATSA |
|---|---|---|---|---|
| *p*-value threshold | Coefficient (95% CI) n = 243.734 | Coefficient (95% CI) n = 1.037 | Coefficient (95% CI) n = 6.221 | Coefficient (95% CI) n = 548 |
| $5x10^{-8}$ | 0.003 (-0.001, 0.007) | -0.033 (-0.093, 0.027) | na | -0.021 (-0.086, 0.045) |
| $1x10^{-5}$ | 0.009 (0.005, 0.012) *** | 0.033 (-0.027, 0.093) | na | -0.006 (-0.088, 0.076) |
| 0.001 | 0.033 (0.029, 0.037) *** | 0.019 (-0.041, 0.079) | 0.033 (0.011, 0.055) ** | 0.015 (-0.071, 0.101) |
| 0.01 | 0.049 (0.045, 0.052) *** | 0.020 (-0.040, 0.080) | 0.044 (0.023, 0.066) *** | -0.023 (-0.095, 0.048) |
| 0.05 | 0.057 (0.053, 0.061) *** | 0.019 (-0.040, 0.078) | 0.047 (0.025, 0.068) *** | -0.011 (-0.079, 0.056) |
| 0.1 | 0.059 (0.056, 0.063) *** | 0.016 (-0.043, 0.075) | 0.048 (0.026, 0.069) *** | 0.000 (-0.073, 0.074) |
| 0.2 | na | na | 0.046 (0.024, 0.068) *** | na |
| 0.3 | na | na | 0.045 (0.023, 0.066) *** | na |
| 0.5 | 0.060 (0.057, 0.064) *** | 0.022 (-0.037, 0.082) | 0.044 (0.022, 0.065) *** | 0.016 (-0.053, 0.086) |
| 1 | 0.061 (0.057, 0.064) *** | 0.015 (-0.043, 0.075) | 0.045 (0.023, 0.066) *** | 0.018 (-0.054, 0.090) |

Coefficients are standardized; effects of a SD increase in $PRS_N$ on a SD increase in FI scores

*$p<0.05$; **$p<0.01$; ***$p<0.001$
